# Restoring striatal WAVE-1 improves maze exploration performance of GluN1 knockdown mice

**DOI:** 10.1101/341479

**Authors:** Yuxiao Chen, Marija Milenkovic, Ali Salahpour, Scott H. Soderling, Amy J. Ramsey

## Abstract

NMDA receptors are important for cognition and are implicated in neuropsychiatric disorders. GluN1 knockdown (GluN1KD) mice have reduced NMDA receptor levels, striatal spine density deficits, and cognitive impairments. However, how NMDA depletion leads to these effects is unclear. Since Rho GTPases are known to regulate spine density and cognition, we examined the levels of RhoA, Rac1, and Cdc42 signaling proteins. Striatal Rac1-pathway components are reduced in GluN1KD mice, with Rac1 and WAVE-1 deficits at 6 and 12 weeks of age. Concurrently, medium spiny neuron (MSN) spine density deficits are present in mice at these ages. To determine whether WAVE-1 deficits were causal or compensatory in relation to these phenotypes, we intercrossed GluN1KD mice with WAVE-1 overexpressing (WAVE-Tg) mice to restore WAVE-1 levels. GluN1KD-WAVE-Tg hybrids showed rescue of striatal WAVE-1 protein levels and MSN spine density, as well as selective behavioral rescue in the Y-maze and 8-arm radial maze tests. GluN1KD-WAVE-Tg mice expressed normalized WAVE-1 protein levels in the hippocampus, yet spine density of hippocampal CA1 pyramidal neurons was not significantly altered. Our data suggest a nuanced role for WAVE-1 effects on cognition and a delineation of specific cognitive domains served by the striatum. Rescue of striatal WAVE-1 and MSN spine density may be significant for goal-directed exploration and associated long-term memory in mice.

## Introduction

The N-methyl-D-aspartate (NMDA) receptor (NMDAR) is a glutamate- and glycine-gated ion channel composed of four subunits: two obligatory GluN1 subunits and a combination of GluN2 or GluN3 family subunits (1). De novo point mutations in GluN1 or GluN2 subunits have been identified in patients with schizophrenia, autism, intellectual disability, and epilepsy (2). Cognitive impairments are prominent features of these neurological disorders. The consequences of NMDAR hypofunction can be studied using GluN1 knock-down (GluN1KD) mice, which have a global genetic reduction of NMDARs by more than 90% (3). Behavioral and cellular phenotypes of GluN1KD mice have been used to model aspects of both schizophrenia and autism (3–6).

At the cellular level, GluN1KD mice have decreased spine density in the striatum that is detected at six weeks of age but not at two weeks. This is temporally correlated with synaptic deficits of the Disrupted in Schizophrenia-1 (DISC1) protein and the onset of cognitive impairments (7,8). DISC1 reductions may mediate dendritic spine deficits since DISC1 is known to regulate Rac1 and neurite growth downstream of NMDARs (9,10). Dendritic spines represent important neuronal connections whose alterations in morphology and number are associated with behavioral changes (11,12). Thus, we hypothesized that the age-dependent reduction of synaptic DISC1 and spine density in GluN1KD mice contributed to cognitive behavioral abnormalities through changes in Rho GTPase signaling (7).

Studies of Rho GTPases (Rac1, RhoA, and Cdc42 in particular) suggest that they work with NMDARs to regulate synaptic plasticity and cognition (13,14). For example, the activity of RhoA and Cdc42 increases in an NMDAR-dependent manner during long-term potentiation (15). Rho GTPases and their downstream signaling molecules also influence cognition at a macroscopic level. Learning and memory deficits result with RhoA inhibition, Rac1 depletion from forebrain excitatory neurons, or with genetic disruption of a downstream effector of Rac1, the Wiskott-Aldrich syndrome protein-family verprolin-homologous protein family member 1 (WAVE-1, also called Scar1) (16–18). Disruptions in Rho GTPase signaling are also linked to neuropsychiatric disorders, including schizophrenia (19,20). We therefore hypothesized that NMDAR hypofunction in GluN1KD mice results in dysregulated Rho GTPase signaling and altered synaptic connectivity that in turn contribute to abnormal GluN1KD mouse behaviors.

To test our hypothesis, we studied the relationship between changes of Rho GTPase pathways proteins, dendritic spines, and behavioral test performance in mice with NMDAR deficiency. We report that GluN1KD mice showed an age-dependent decrease of dendritic spine density in striatal medium spiny neurons (MSNs). Striatal Rac1 signaling was also changed in 3, 6, and 12 week old GluN1KD mice – consistently showing lower Rac1 and WAVE-1 levels in older mice. We tested whether WAVE-1 over expressing (*Wasf1* – Entrez Gene ID 8936 – BAC transgenic, WAVE-Tg) mice could rescue the cellular or behavioral phenotypes of GluN1KD mice. GluN1KD-WAVE-Tg compound transgenic mice had improved striatal WAVE-1 levels and MSN spine density. They also showed improved performance in the Y-maze and 8-arm radial maze tests, but not in other behavioral tests in our study. Further assessments of WAVE-1 and dendritic spine changes at the hippocampus found no significant changes resulting from the knock-down of GluN1 subunits or the addition of a *Wasf1* transgene. Our results point to a nuanced role for WAVE-1 in synaptic plasticity and cognition, particularly in the striatum, regarding maze exploration performance.

## Methods

### Animals

WT, WAVE-Tg, GluN1KD, and GluN1KD-WAVE mice aged 3, 6, or 12-14 weeks were used in this study. All experiments used only adult mice aged 12-14 weeks except for those explicitly involving and comparing 3 and 6-week old mice. Male and female mice were used for all experiments except for dendritic spine studies, RNAscope analysis, and hippocampal WAVE-1 western blots of all four genotypes, which used males. The generation of GluN1KD mice was previously described (3), in which a neomycin cassette was inserted into intron 19 of *Grin1* (Entrez Gene ID 14810). The mouse line was backcrossed for more than 20 generations onto two genetic backgrounds, C57B1/6J and 129X1/SvJ. Mice used for comparisons between GluN1KD and WT littermates were generated from intercross breeding of C57B1/6J *Grin1*+/− and 129X1/SvJ *Grin1*+/− heterozygotes to produce F1 progeny. This strategy was chosen in accordance with the recommendations of the Banbury Conference to minimize the potential confound of homozygous mutations in parent strains (21).

WAVE-Tg mice were constructed by the pronuclear injection of BAC DNA containing the human *Wasf1* locus into fertilized eggs of C57B1/6J mice. Mice used to compare between WT, WAVE-Tg, GluN1KD and GluN1KD-WAVE mice were generated by intercross breeding between C57B1/6J *Grin1*+/− WAVE-Tg mice and 129X1/SvJ *Grin1*+/− non-WAVE-Tg mice. All F1 mice were genotyped by polymerase chain reaction analysis of tail sample DNA. The primers used to amplify the GluN1KD *Grin1*– allele were: 5′ AAG CGA TTA GAC AAC TAA GGG T 3′ and 5′ GCT TCC TCG TGC TTT ACG GTA T 3′. The primers used to amplify the WT *Grin1* allele were: 5′ TGA GGG GAA GCT CTT CCT GT 3′ and 5′ AAG CGA TTA GAC AAC TAA GGG T 3′. For WAVE-1 genotyping, the *Wasf1*-targeting primers 5′ CAA CTC ATT GCA AGA ACG TGT GGA C 3′ and 5′ AAT AAA AAA ATT AGC CAG GCG TGG TG 3′ were used. All animal housing and experimentation conditions and protocols were in accordance with institutional (University of Toronto Faculty of Medicine and Pharmacy Animal Care Committee, approved protocol #20011988) and federal (Canadian Council on Animal Care) guidelines. Behavioral tests were performed during the light cycle, between 9am and 6pm. Mice were naïve to all behavioral tests before use in this study. To minimize animal suffering, mouse brains were collected after quick and precise cervical dislocation for biochemical measures, and after terminal anesthesia with tribromoethanol for spine density measures.

### Dendritic spine analysis

Dendritic spine density and morphology were assessed via DiOLISTIC labeling and confocal microscopy as detailed previously (22,23). Mice were deeply anesthetized with tribromoethanol and perfused transcardially with phosphate buffered saline (PBS) followed by a solution of 4% paraformaldehyde (PFA) in PBS over 10 minutes. Brains were then collected and submerged in 4% PFA for one hour before being stored in PBS at 4°C. Within 48 hours, perfused brains were sliced coronally at a thickness of 100 or 150µm. Slices of the striatum and hippocampus were collected and their neurons were randomly labeled by DiI (1-1′-Dioctadecyl-3,3,3′,3′-tetramethylindocarbocyanine perchlorate) labeling. Slices were impregnated with DiI coated tungsten or gold beads using a Helios gene gun (Bio-Rad, CA, USA).

Z-stack images with steps of 0.5-0.9 μm at 60X magnification were collected via an IX81 confocal microscope and Fluoview FV 1000 software (Olympus, Tokyo, Japan). Images comparing WT and GluN1KD mice at 3, 6, and 12 weeks of age were analyzed for changes in spine density with NIS-Elements Basic Research (Version 3.10, Nikon, Tokyo, Japan). Images comparing spine density between all four genotypes were analyzed with ImageJ (Version 1.50; (24) from the National Institutes of Health (NIH, MD, USA) using the NeuronJ (Version 1.4.3) plug-in (25). Striatal MSN dendrites were assessed randomly in the initial comparison between WT and GluN1KD mice using 100 µm thick brain slices. Later striatal spine assessments comparing WT, WAVE-Tg, GluN1KD and GluN1KD-WAVE mice were focused on 20 µm dendrite segments starting approximately 20 µm away from the soma and used 150 µm thick slices. Hippocampal slices 150 µm thick were analyzed at 20 µm segments of secondary and tertiary dendrites from the basolateral and apical sides of CA1 pyramidal neurons. Basolateral dendrites were assessed starting at 30-45 µm away from the soma while apical dendrites were assessed near the initial branching points of secondary or tertiary dendrites closest to the soma, which varied in distance from soma. All dendritic segments selected for analysis avoided branch points. Two to six z-stack image compositions were used for each brain region and mouse assessed. Spine density analyses were performed blind to genotype.

### Western blotting

The striatum and hippocampus of test mice were dissected to prepare protein extracts. Each brain region was homogenized in 400 µl of PHEM buffer (0.5% TritonX 100, 60mM PIPES, 25mM HEPES, 10mM EGTA, and 2mM MgCl_2_) with protease and phosphatase inhibitors (1.5 µg/ml aprotinin, 10 µg/ml leupeptin, 10 µg/ml pepstatin A, 0.1mg/ml benzamidine, 0.25mM PMSF, 5mM Na orthovanadate, 10mM NaF, 2.5mM Na pyrophosphate, 1mM β-glycerophosphate). Loading samples were prepared from protein homogenates with standard sample buffer and 5% β-mercaptoethanol, then heated at 100°C for 5 minutes. Depending on the protein of interest, 15-25 μg of protein for each sample were electrophoresed in 7.5-12% bis-acrylamide gels and transferred onto polyvinylidene difluoride membranes, both steps at 80-100V. Blots were blocked with 5% milk or bovine serum albumin in tris-buffered saline and Tween 20, according to primary antibody supplier instructions. Blots were incubated with fluorescent secondary antibodies and imaged with the Odyssey Infrared Imaging System (LI-COR, NE, USA). Densitometry was performed using Odyssey (Version 3, LI-COR), Image Studio (Version 5, LI-COR), or ImageJ (Version 1.45, NIH). Primary antibodies used were as follows: α-WAVE-1 (1:1000-2000, catalog number 75-048, NeuroMab, CA, USA), α-Rac1 (1:500, catalog number ARC03, Cytoskeleton, CO, USA), and α-GAPDH (1:4000-5000, catalog number G8795, Sigma, MO, USA). Secondary antibodies were as follows: donkey α-mouse IgG-800 (1:5000, catalog number 610-731-002, Rockland, PA, USA), goat α-rabbit IgG-680 (1:5000, catalog number A21076, Invitrogen, CA, USA), donkey α-mouse IgG-680 (1:15000, catalog number 926-68072, LI-COR, NE, USA) and goat α-rabbit IgG-800 (1:15000, catalog number 926-32211, LI-COR). Blots were normalized to loading control protein (GAPDH) bands or to REVERT total protein stain (LI-COR, NE, USA) before analysis.

### Fluorescent in situ hybridization via RNAscope

*Wasf1* expression in adult male mice of all four genotypes was visualized by RNAscope, a chromogenic *in situ* hybridization technique, following the RNAscope Multiplex Fluorescent Reagent Kit v2 from ACD Bio (CA, USA). Fresh-frozen brains were sliced sagittally (lateral 1.2 mm Bregma) at a thickness of 20µm and mounted onto charged slides. Slices were then fixed by submersion into 4% PFA in 1X PBS and dehydrated by serial submersion in 50%, 70%, and 100% ethanol. Slices were then incubated in hydrogen peroxide before being treated with RNAscope Protease IV. The multiplex fluorescent assay was then performed: slices were hybridized to negative control (reference ID 320871, ACD Bio), positive control (ref ID 320881), or both human (ref ID 533691) and mouse (ref ID 533701) specific *Wasf1* probes; amplification reagents were then used to incubate the slides to increase signal strength; finally, fluorescent signals were developed using TSA Plus cyanine 3 (PerkinElmer, Llantrisant, UK) for channel 1 and cyanine 5 (also PerkinElmer) for channel 2 using channel-specific horseradish peroxidases. Vectashield with DAPI (Vector Laboratories, CA, USA) was then applied to the slices before glass coverslip placement. Slides were dried overnight and then imaged with an Axio Scan.Z1 slide scanner (Zeiss, Oberkochen, DEU) at 20X magnification.

### Behavioral tests

#### Y-maze spontaneous alternation test

The Y-maze spontaneous alternation test was performed as an assessment of cognitive function of test mice, as described in previous studies (8,26). For 8 minutes, mice were allowed to freely explore a Y-maze consisting of three identical arms with the dimensions 38 × 7.6 × 12.7 cm that met at the center separated by an angle of 120° between each pair of arms (San Diego Instruments, CA, USA). Mice were tracked once they were placed at the end of one arm with Biobserve Viewer (Version 2; St. Augustin, Germany). Zones were digitally defined for each arm starting at 5 cm away from the center of the maze. Spontaneous 3-arm alternations were defined as a mouse executing consecutive entries into each of the three zones without repeat entries into any one zone. The percent of spontaneous three-arm alteration zone entries relative to all Y-maze zone entries after the first two was taken as the Y-maze performance score.

#### 8-arm radial maze test

WT, WAVE-Tg, GluN1KD, and GluN1KD-WAVE mice were also assessed for their cognitive function based on the 8-arm radial maze as described (Dzirasa et al., 2009). The maze consisted of 8 arms with the dimensions 22.86 × 7.62 × 15.24 cm that met together as sides of an octagon, which served as the maze center (San Diego Instruments, CA, USA). Mice were food restricted to 90% of their free-feeding body weight. After habituation to the maze center for one week, the arms were baited with a piece of cherrio cereal and mice explored the arena for 5 minutes or until all 8 arms were explored. Mice were tested daily for four days, and the performance of the four days was averaged for each week (each trial block). Cognition was assessed by measures of working memory error (WME, the number of entries into already-visited arms) and by measures of entries-to-repeat (ETR, the number of arms visited before the mouse makes a repeat entry into any arm) recorded by Biobserve Viewer (Version 3).

#### Puzzle box test

Problem-solving, short-term memory and long-term memory were assessed by the puzzle box test as described previously (8,27). The puzzle box has an arena with two adjacent compartments measuring 58 × 28 × 27.5cm and 14 × 28 × 27.5cm that are separated by one of two removable dividers: one has an open door allowing free passage while the other leaves only an underpass connection. To start each trial, mice were placed in the larger arena compartment under bright-light conditions facing away from the divider. As the other compartment beyond the divider has a roof cover providing dim-light conditions, mice are motivated to enter it. The time that it took the mouse to completely enter the smaller area was manually recorded. There were 9 trials spread out to 3 per day with new obstacles introduced during the second trial of each day to block the path between the two arena compartments. Replacement of the open door divider with the door-less divider was introduced in trial 2 and was used for every subsequent trial. The underground pass was filled with cage bedding in trials 5-7, then instead with a cardboard plug in trials 8-9. 5 minutes were given for each mouse to reach the dim-light area during each trial and 2 minutes separated trials for the same mouse on the same day. Mice that failed the first trial, a training trial, were excluded from the test.

#### Open field test

The protocol for the open field test of locomotor activity has been described previously (3,28). Test mice were placed in a novel environment, clear Plexiglas chambers measuring 20 × 20 × 45cm, for 2 hours. Their locomotor activity was recorded and tracked by digital activity monitors from Omnitech Electronics (OH, USA) via infrared beam sensors. Distance travelled was analyzed in five-minute bins to assess locomotor activity.

#### Social approach behavior test

Test mice were also assessed for their social cognition via approach behavior towards novel age- and sex-matched mice, which served as social stimuli. A social approach behavior test was performed as previously described (29,30). Test mice were placed in a white Plexiglas arena (62 × 40.5 × 23cm). Their subsequent interactions with an empty inverted wire cup and another containing the social stimulus mouse were recorded with a video camera for 10 minutes. Biobserve Viewer (Version 2) was used to track center of body mass and quantify the amount of time mice spent in 5cm circular zones surrounding each cup. The amount of time spent by test mice in both the social stimulus and nonsocial zones were taken as a measure of social cognition and a related novelty control measure, respectively.

### Statistical analysis

All data were graphed and analyzed using Prism (Version 6.01, GraphPad Software, CA, UAS), except for two-way ANOVAs with repeated measures, which were analyzed using SPSS (Version 20.0.0, IBM, NY, USA). Assumptions of equal variance were tested for all independent measures tests while assumptions of sphericity were tested for all repeated-measures tests. Comparisons between WT and GluN1KD mice were assessed with independent, two-tailed t-tests while those between WT, WAVE-Tg, GluN1KD, and GluN1KD-WAVE mice were assessed with one-way ANOVAs except in cases with repeated measurements over time or in cases where equal variance could not be assumed. Comparisons of repeated-measures data, including locomotor activity and 8-arm radial maze performance, were completed using two-way ANOVAs with repeated measures. Bonferroni corrections were used after ANOVAs for *post hoc* comparisons between all groups. Data with unequal variance were compared with Kruskal-Wallis tests followed by *post hoc* Dunn’s multiple comparisons tests. Behavioral test results were analyzed for sex differences. A cut-off of *p*-value of 0.05 was chosen for statistical significance. Sample sizes are indicated within each figure for all experiments. Data are expressed as mean ± standard error of the mean. Where mentioned, *post hoc* power analyses and sample size calculations were performed with G*Power (Version 3.1.9.2; (31,32) from the Heinrich Heine University Düsseldorf).

## Results

### GluN1KD mice have age-dependent deficits in striatal MSN dendritic spine density and aberrant levels of Rac1 signaling components

We previously reported that striatal spine density of GluN1KD mice was normal at two weeks of age but was decreased at six weeks of age relative to WT littermates (7). We therefore assessed spine density at 3, 6, and 12 weeks of age to further study the onset and progression of developmental spine loss. In the striatum, GluN1KD mice have spine deficits at 6 and 12 weeks of age, but not at 3 weeks (Figure 1). GluN1KD mice had a 11% reduction in spine density at 6 weeks (WT: 138 ± 2 spines/100 μm, GluN1KD: 124 ± 2 spines/100 μm) (independent, two-tailed *t_(4)_* = 5.55, *p* < 0.01). At 12 weeks of age GluN1KD mice had a 16% reduction in spines (WT: 136 ± 2 spines/100 μm, GluN1KD: 114 ± 1 spines/100 μm) (*t_(4)_* = 9.74, *p* < 0.01). Thus, we determined that spine loss first occurs between 3 and 6 weeks of age, and that these reductions are maintained in the adult brain.

**Figure 1.**
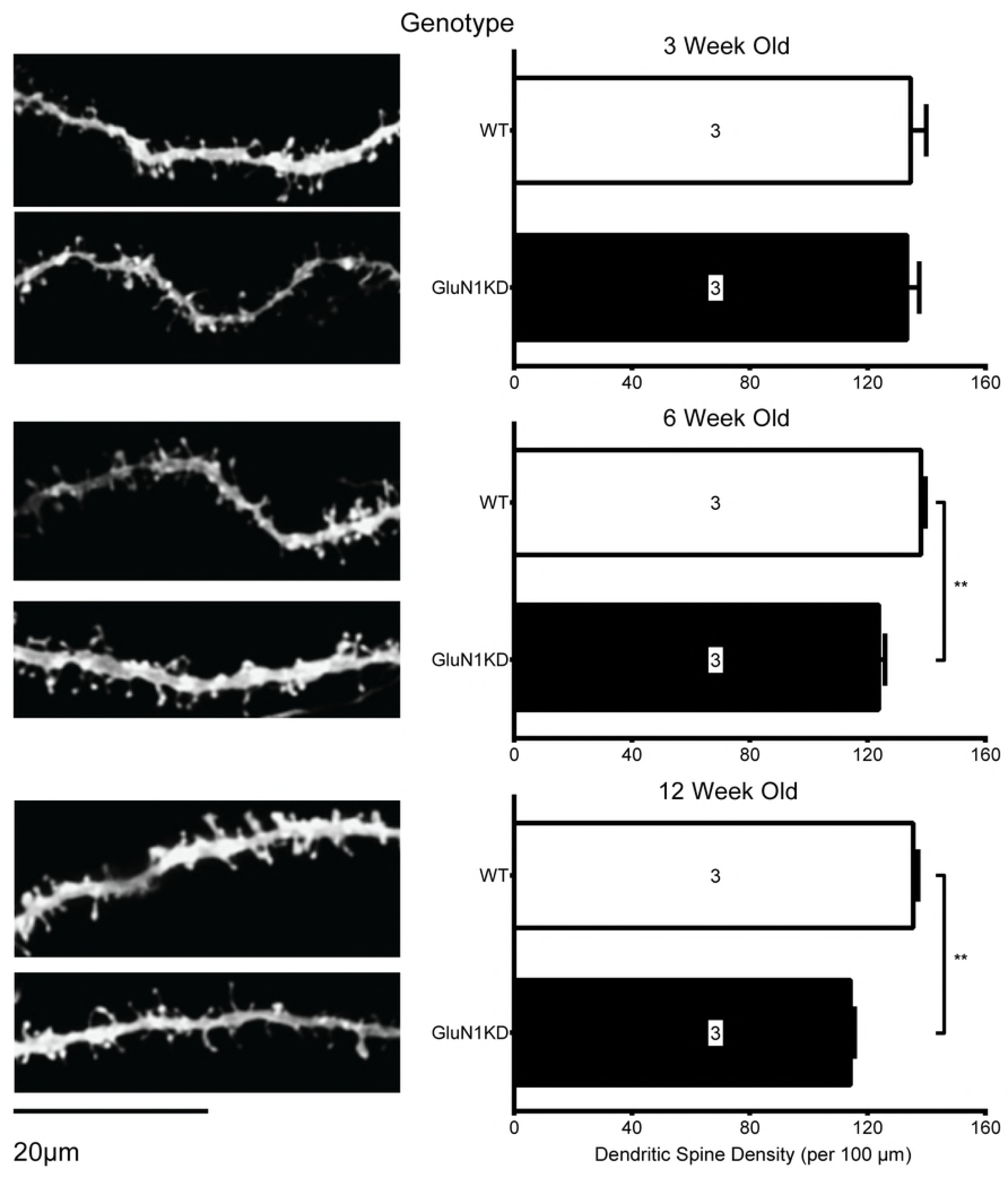
Representative images (left) and quantifications (right) of dendritic spines of striatal MSNs in GluN1KD and WT mice at 3, 6, and 12 weeks of age. GluN1KD mice have significantly reduced spine density compared to WT mice at 6 and 12 weeks of age. Mouse sample sizes are denoted within each bar. 3-7 dendrite sample images were analyzed for each mouse. Scale bar represents 20µm. Data was analyzed by two-tailed, independent t-tests, \*\**p* < 0.01.

We measured the protein levels of key components of Rho GTPase signaling cascades in the striatum of 3, 6, and 12-week-old mice. The proteins assessed were RhoA, Rac1, Cdc42, cortactin, WAVE-1, N-WASP, LIMK1, cofilin, pS3-cofilin, and actin. Of these, significant differences were consistently found for Rac1 and WAVE-1 (Figure 2). GluN1KD mice show an increase in Rac1 at 3 weeks, and reductions in Rac1 at 6 and 12 weeks of age. Expressed as a percentage of WT levels, GluN1KD levels of Rac1 are 126% at 3 weeks, 70% at 6 weeks, and 76% at 12 weeks of age (independent, two-tailed *t_(6)_* = 2.45-6.00, *p* < 0.05 for all three). WAVE-1, a downstream effector of Rac1, is reduced in the GluN1KD striatum at all three ages. Expressed as a percentage of WT, WAVE-1 levels are 75 % at 3 weeks, 63% at 6 weeks, and 80% at 12 weeks of age (*t_(6)_* = 2.60-3.04, *p* ≤ 0.05 for all three). Of the Rho GTPase signaling proteins, the Rac1 pathway proteins are especially altered in GluN1KD mice - specifically Rac1 itself and its downstream effector, WAVE-1. (33,34).

**Figure 2.**
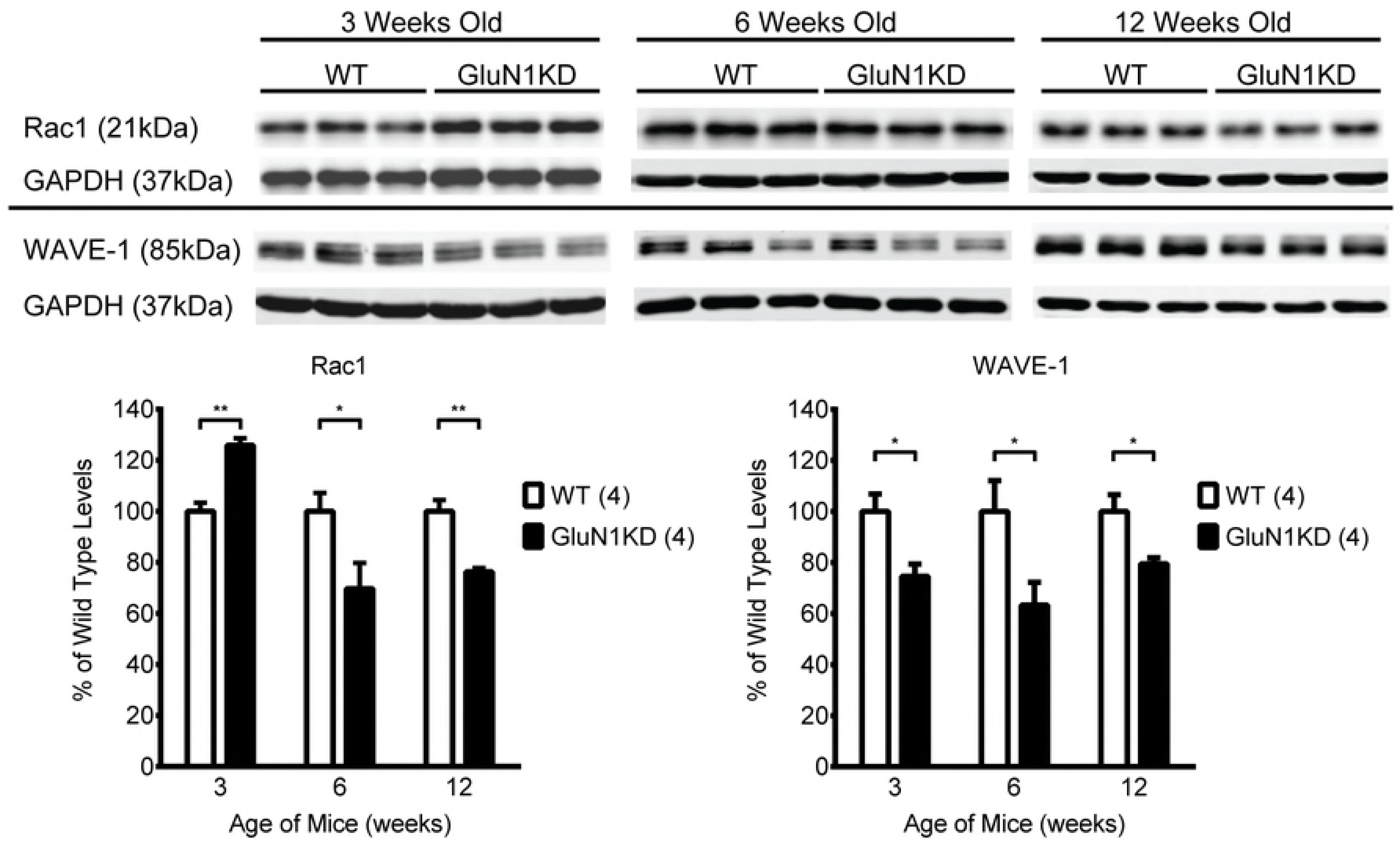
Representative western blots (top) and their quantifications (bottom) assessing differences in Rac1 and WAVE-1 levels at the striatum of GluN1KD mice and WT littermates aged 3, 6, and 12 weeks. In GluN1KD mice, Rac1 was significantly increased at 3 weeks before decreasing to lower than WT levels at 6 and 12 weeks of age. In contrast, WAVE-1 was consistently decreased at all time points. All blots were first normalized to GAPDH loading controls. A sample size of 4 mice per group, denoted in the graph legends beside genotype labels, was used for every genotype and age group. Data was analyzed by two-tailed, independent t-tests, \**p* ≤ 0.05, \*\**p* < 0.01.

### Striatal WAVE-1 levels are increased in WAVE-Tg mice and restored in GluN1KD-WAVE hybrids

In other model systems, reductions in Rac1 and WAVE-1 cause dendritic spine loss (18,35), but chronic increases in Rac1 activity can also cause spine abnormalities (9,36,37). Therefore, we considered the possibilities that reductions in Rac1 and WAVE-1 could either contribute to the observed spine loss or could be compensatory responses to prevent further spine loss. To test these two possibilities, we restored WAVE-1 protein levels in GluN1KD mice by intercross breeding with a transgenic mouse line that overexpresses WAVE-1.

WAVE-1 transgenic mice (WAVE-Tg) were generated by pronuclear injection of a bacterial artificial chromosome bearing the entire the human *Wasf1* genomic sequence with 86.5 kb of upstream and 70.3kb of downstream genomic sequence. *Wasf1* transgenic mice (WAVE-Tg and GluN1KD-WAVE-Tg hybrids, called GluN1KD-WAVE hereafter) show increased WAVE-1 message expression compared to non-transgenic littermates at the striatum (Figure 3) and hippocampus (Figure 4). These increases are specific to the human transgene and similar to endogenous mouse *Wasf1* expression in pattern. We also performed western blotting of striatal protein to measure the relative levels of Rac1 and WAVE-1 in this new line of mice (Figure 5). One-way ANOVA found no significant genotype effect on Rac1 levels (One-way ANOVA *F_3,20_* = 1.514, *p* = 0.24, *β* = 0.66) but shows a significant effect of genotype on levels of WAVE-1 (*F_3,20_* = 33.59, *p* < 0.01). The presence of the WAVE-Tg increases the levels of WAVE-1 in the striatum of both WT and GluN1KD mice. WAVE-1 is 120% of WT levels in WAVE-Tg mice (Bonferroni adjusted *p* < 0.01 vs WT), 69% in GluN1KD mice (*p* < 0.01), and 87% in GluN1KD-WAVE-Tg hybrids (not significantly different vs WT). These results indicate that GluN1KD mice have reduced WAVE-1 levels and that the *Wasf1* transgene is sufficient to rescue striatal WAVE-1 levels in GluN1KD mice.

**Figure 3.**
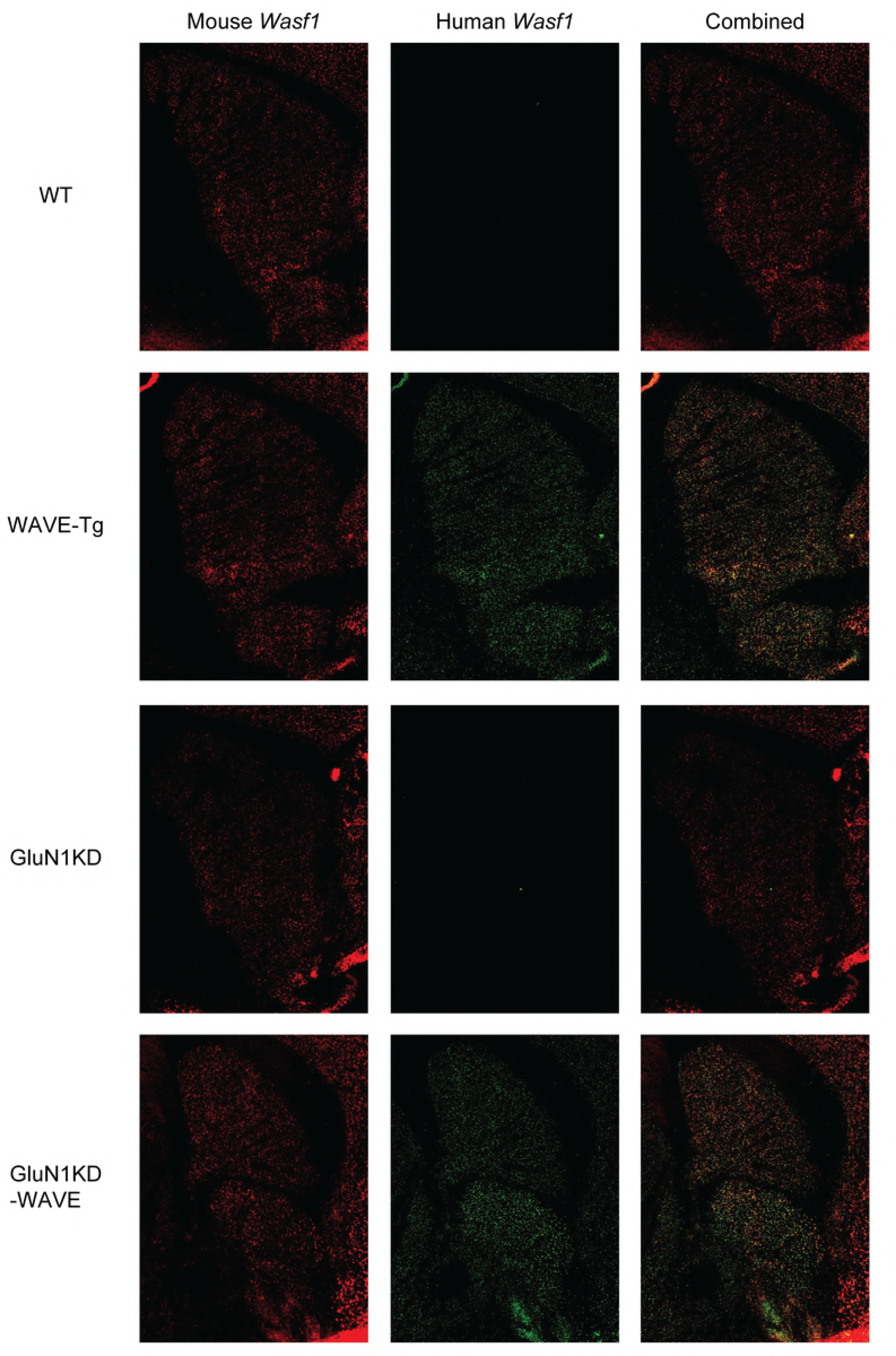
Fluorescent *in situ* hybridization of WT, WAVE-Tg, GluN1KD, and GluN1KD-WAVE striatum. Endogenous, mouse *Wasf1* message is shown in red (left column) while transgenic, human *Wasf1* is shown in green (middle column). Human *Wasf1* was expressed abundantly and specifically in transgenic mice with a pattern of expression similar to mouse *Wasf1*.

**Figure 4.**
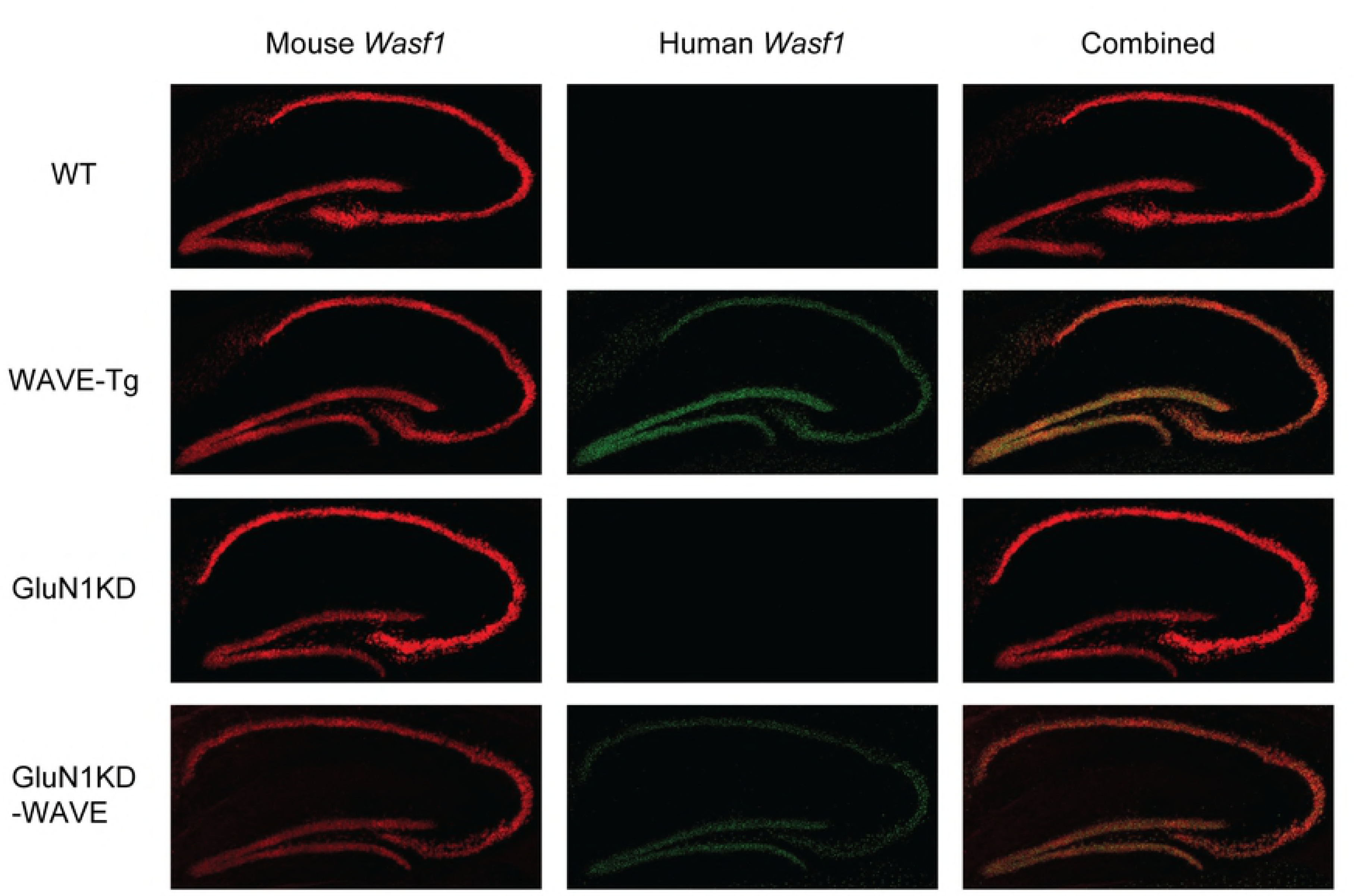
Fluorescent *in situ* hybridization of WT, WAVE-Tg, GluN1KD, and GluN1KD-WAVE hippocampus. Endogenous, mouse *Wasf1* message is shown in red (left column) while transgenic, human *Wasf1* is shown in green (middle column). Human *Wasf1* was expressed abundantly and specifically in transgenic mice with a pattern of expression similar to mouse *Wasf1*.

**Figure 5.**
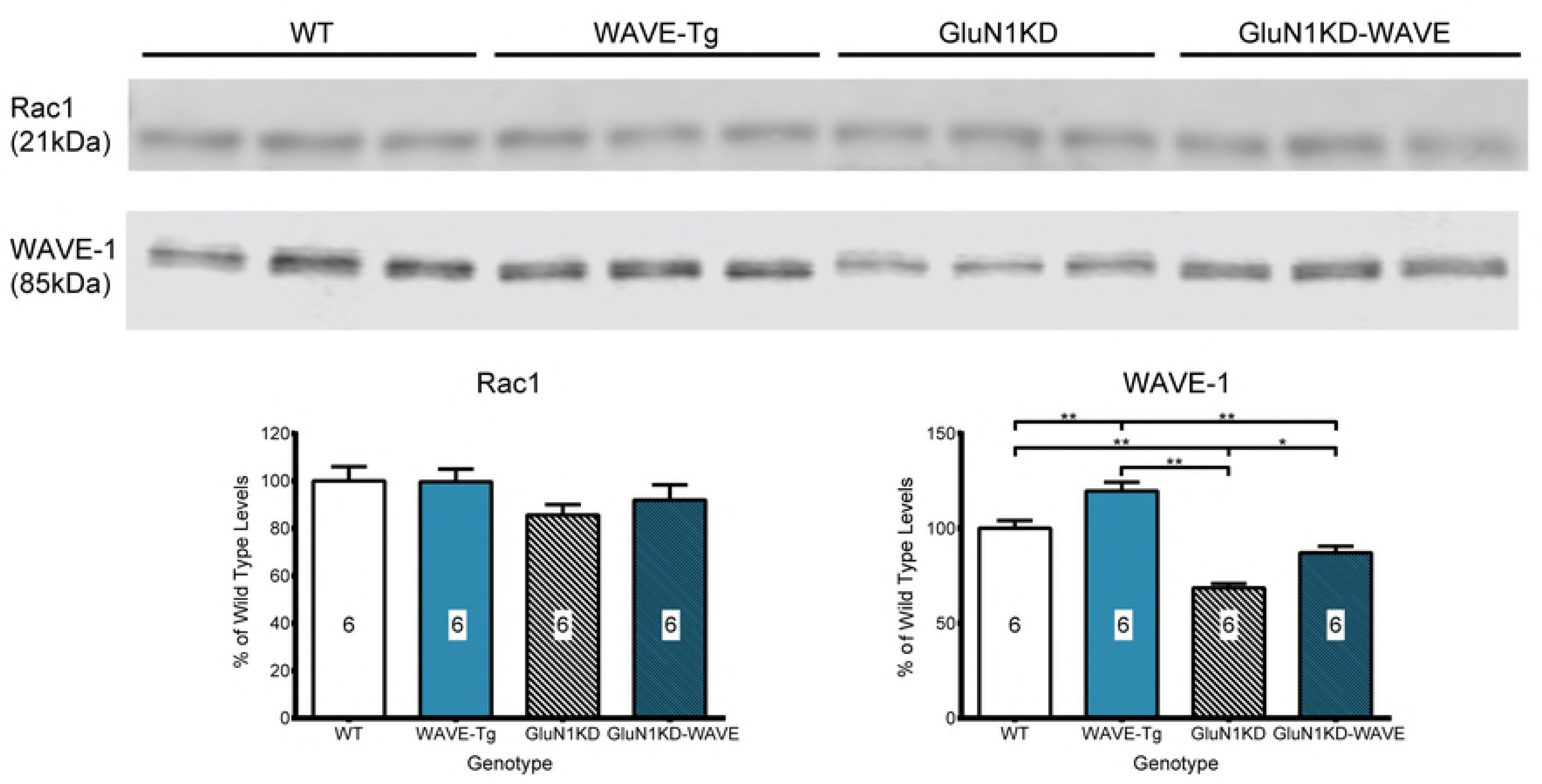
Representative western blots (top) and their quantifications (bottom) assessing differences in striatal levels of Rac1 and WAVE-1 of adult WT, WAVE-Tg, GluN1KD, and GluN1KD-WAVE mice. WAVE-1 was confirmed to be decreased in GluN1KD mice, increased in WAVE-Tg mice, and attenuated towards WT levels in GluN1KD-WAVE mice. Rac1 was not found to be significantly different between genotypes. A sample size of 6 mice was used for every genotype. Data was analyzed by one-way ANOVAs followed by Bonferroni *post hoc* comparisons for all pairings, \**p* < 0.05, \*\**p* < 0.01.

### Striatal spine density deficits are attenuated in GluN1KD-WAVE hybrids

As we have rescued WAVE-1 expression in GluN1KD-WAVE mice, we next asked whether this attenuates the synaptic deficits of GluN1KD mice in striatal MSNs. We therefore assessed the dendritic spine density of adult WT, WAVE-Tg, GluN1KD, and GluN1KD-WAVE mice. Representative dendrite images and spine density quantifications are shown in Figure 6a. One-way ANOVA reported an effect of genotype on spine density that trended towards significance (*F_3,32_* = 2.79, *p* = 0.06). GluN1KD mice had a spine density of 145 ± 9 spines/100 μm, lower than WT levels of 185 ± 11 spines/100 μm, also trending towards significance (Bonferroni adjusted *p* = 0.05). In contrast, GluN1KD-WAVE mice had a more WT-like spine density of 172 ± 6 spines/100 μm (*p* > 0.99 vs WT). These results point to the improved WAVE-1 levels of GluN1KD-WAVE mice having biological significance at the synapse by increasing spine density. They also led us to test whether GluN1KD-WAVE mice have improved behavioral test performance, which would more definitively assess the biological significance of restoring WAVE-1 in NMDAR-deficient mice.

**Figure 6.**
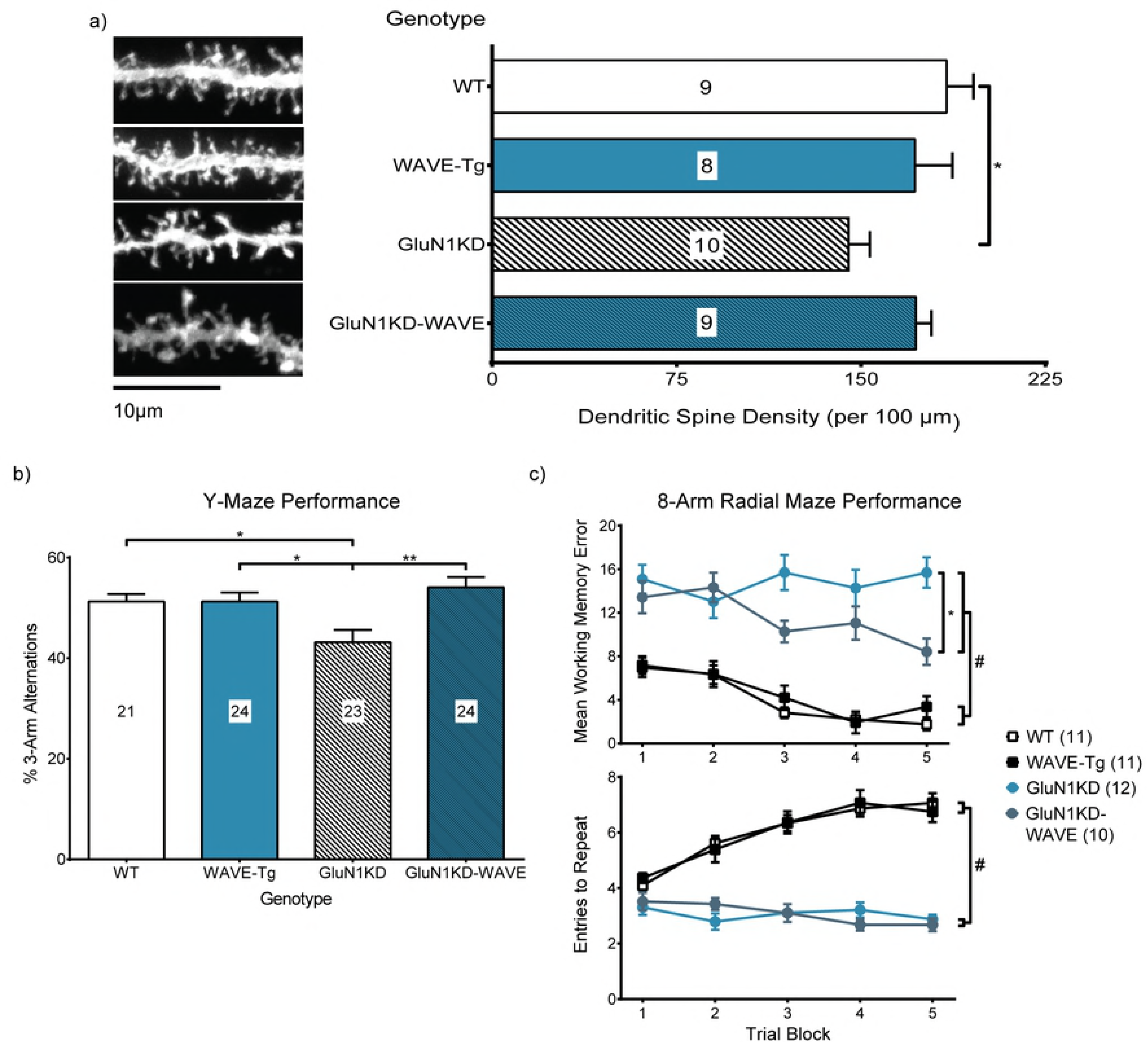
Striatal MSN spine density (a), Y-maze performance (b), and 8-arm radial maze performance (c) are deficient in GluN1KD mice and improved in GluN1KD-WAVE mice. a) Representative images of MSN dendrites (left) and spine density quantifications (right) show lower spine density in GluN1KD mice compared to WT littermates but not in GluN1KD-WAVE hybrids. Mouse sample sizes are denoted within each bar. 3-6 dendrite sample images were analyzed for each mouse. Scale bar represents 20µm. Data was analyzed by one-way ANOVA followed by Bonferroni *post hoc* comparisons for all pairings, * *p* = 0.05. b) GluN1KD, but not GluN1KD-WAVE, mice had significantly lower % 3-arm alternation scores compared to WT and WAVE-Tg mice. GluN1KD-WAVE mice also had a significantly higher score compared to GluN1KD mice. Data was analyzed by one-way ANOVA followed by Bonferroni *post hoc* comparisons for all pairings, * *p* < 0.05, ** *p* < 0.01. Sample sizes are indicated by numbers in each bar. c) Mean WME (top) and ETR (bottom) scores from the 8-arm radial maze test show different effects of intercrossing GluN1KD and WAVE-Tg mice. GluN1KD and GluN1KD-WAVE mice performed worse on both measures compared to WT and WAVE-Tg mice, but GluN1KD-WAVE mice also had significantly less WMEs over the course of the experiment compared to GluN1KD mice. Two-way repeated measures ANOVAs reported significant effects of genotype on both WME and ETR (*p* < 0.01). # Bonferroni *post hoc* analysis reported an adjusted *p* < 0.01 for each comparison between a GluN1KD and a non-GluN1KD genotype. * *p* = 0.02 for GluN1KD-WAVE mice WMEs compared to GluN1KD mice. In the context of 8-arm radial maze WMEs, intercrossing GluN1KD and WAVE-Tg mice resulted in a partial rescue. No sex differences were observed in these behavior tests. Sample sizes are indicated next to their respective genotypes in the legend.

### Maze navigation test performance is improved in GluN1KD-WAVE mice

#### Y-maze spontaneous alternation test

The Y-maze spontaneous alternation test was performed to assess the cognitive function of test mice. One-way ANOVA revealed a significant effect of genotype on Y-maze spontaneous alternation (*F_3,88_* = 5.70, *p* < 0.01). Shown in Figure 6b, GluN1KD mice perform fewer spontaneous 3-arm alternations (43.2 ± 2.4 %) than WT (51.3 ± 1.5 %, Bonferroni adjusted *p* = 0.04) and WAVE-Tg mice (51.3 ± 1.8 %, *p* = 0.03) while GluN1KD-WAVE hybrids do not (54.1 ± 2.0 %, *p* = 0.01 vs GluN1KD, *p* > 0.99 vs WT and WAVE-Tg). No sex differences were observed in this behavior test. In the context of Y-maze exploration performance, increasing WAVE-1 levels improves cognitive function.

#### 8-arm radial maze test

The 8-arm radial maze test was performed to confirm and complement our Y-maze test. Both working memory errors (WME) and entries-to-repeat (ETR) were recorded as measures of cognitive function (Figure 6c). Two-way repeated measures ANOVAs report significant effects of genotype on WME and ETR (*F_3,40_* = 53.03, *p* < 0.01; *F_3,40_* = 66.80, *p* < 0.01; respectively). For WME, GluN1KD mice make significantly more repeat-arm entries over the 5 trial blocks compared to WT (Bonferroni adjusted *p* < 0.01), WAVE-Tg (*p* < 0.01), and GluN1KD-WAVE mice (*p* = 0.02). This test reveals an intermediate rescue of working memory, since GluN1KD-WAVE mice are also significantly worse compared to WT (*p* < 0.01) and WAVE-Tg mice (*p* < 0.01). Furthermore, both GluN1KD and GluN1KD-WAVE mice rapidly re-enter an already-explored arm, resulting in lower ETR scores, compared to WT and WAVE-Tg mice (Bonferroni adjusted *p* < 0.01 for each comparison between a GluN1KD and a non-GluN1KD genotype). GluN1KD-WAVE mice are not significantly different compared to GluN1KD mice for ETR scores. No sex differences were observed in this behavior test. When assessed by the 8-arm radial maze test, intercrossing GluN1KD and WAVE-Tg mice results in a selective and partial improvement of working memory.

### GluN1KD-WAVE mice do not show significant improvements in other tested behaviors

#### Puzzle box test

The puzzle box test was also used to compare the cognitive function of GluN1KD-WAVE mice with WT, WAVE-Tg, and GluN1KD littermates. Kruskal-Wallis tests for puzzle box solution times of each trial, which involved different aspects of cognition between them, consistently reveal significant effects of genotype (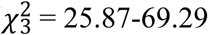, *N*= 94, *p* < 0.01 for all trials). As depicted in Figure 7a, GluN1KD and GluN1KD-WAVE mice take longer to complete each trial compared to WT and WAVE-Tg mice. In trials 5-9, GluN1KD and GluN1KD-WAVE mice often fail to complete the task, requiring the maximum 5-minute exploration time. Dunn’s *post hoc* multiple comparison tests for each trial reveal significantly different rankings (*p* ≤ 0.015) for each comparison between a GluN1KD genotype and a non-GluN1KD genotype. GluN1KD and GluN1KD-WAVE mice were not significantly different. No sex differences were observed in this behavior test. Intercrossing GluN1KD mice with WAVE-Tg mice does not significantly improve cognition in the context of puzzle box test performance.

**Figure 7.**
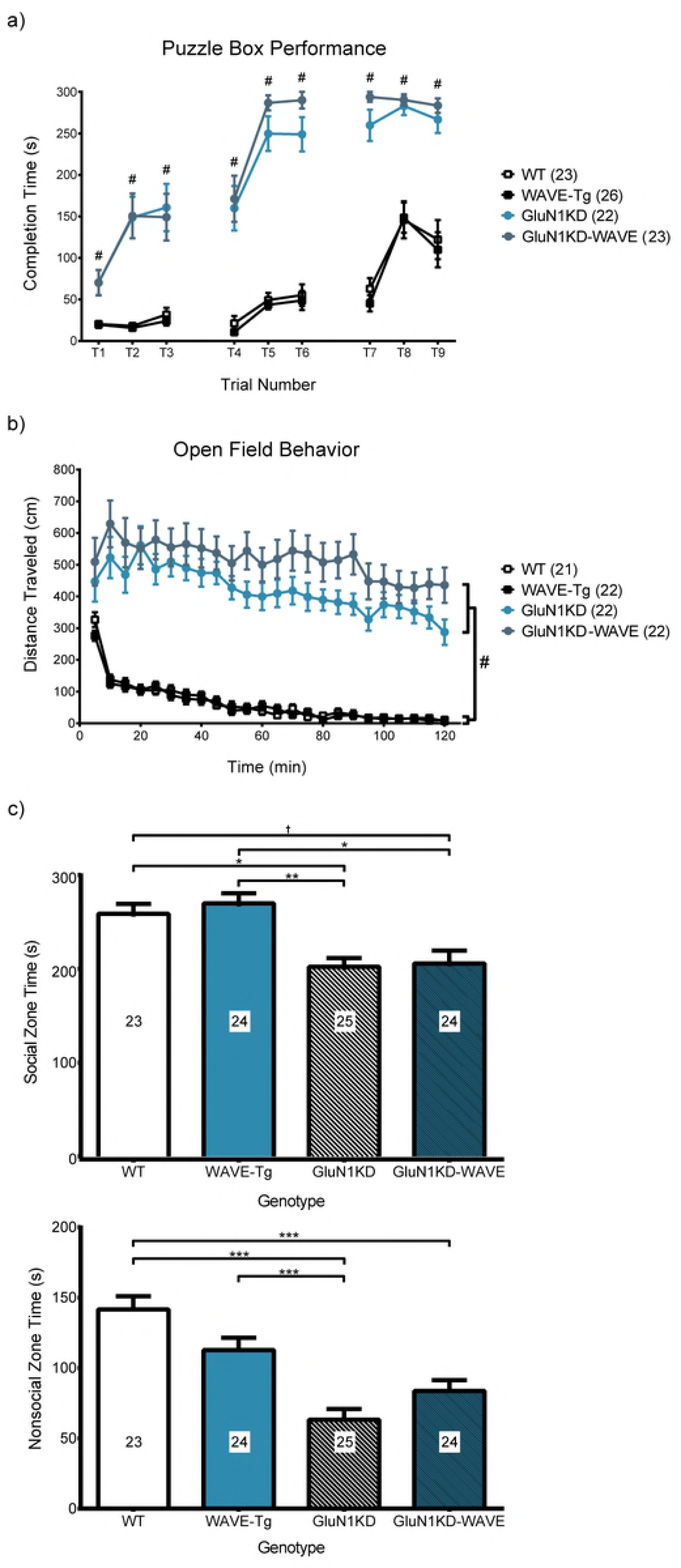
Performance of WT, WAVE-Tg, GluN1KD and GluN1KD-WAVE mice in the (a) puzzle box, (b) open field and (c) social approach behavior tests. **a)** Mice with GluN1KD genotypes took significantly longer to complete puzzle box trials compared to non-GluN1KD mice. Data for each trial was analyzed by independent sample Kruskal-Wallis tests followed by *post hoc* Dunn’s multiple comparisons. # indicates a significant effect of genotype on test completion speed rankings with *p* ≤ 0.015 for each comparison between a GluN1KD genotype and a non-GluN1KD genotype for each trial. GluN1KD and GluN1KD-WAVE mice did not differ significantly in any of the puzzle box trials. Sample sizes are indicated next to their respective genotypes in the graph legend. **b)** Distance traveled over time was recorded in the open field test. Mice of GluN1KD genotypes travelled greater distances and did not display habituation compared to non-GluN1KD littermates. Two-way repeated measures ANOVAs reported a significant main effect of genotype (*p* < 0.01). # Bonferroni *post hoc* analyses reported *p* < 0.01 for each comparison between a GluN1KD and a non-GluN1KD genotype. No significant differences were found between GluN1KD and GluN1KD-WAVE mice. Sample sizes are next to each genotype in the graph legend. **c)** GluN1KD mice spent less time in the social zone, which contained a social stimulus mouse, compared to non-GluN1KD mice. GluN1KD-WAVE mice scored significantly lower compared to WAVE-Tg mice and trended towards significance compared to WT mice († *p* = 0.15). WT mice also spent more time in the nonsocial zone compared to both GluN1KD and GluN1KD-WAVE mice. No significance was found comparing GluN1KD to GluN1KD-WAVE mice. No sex differences were observed in these behavior tests. Data was analyzed by one-way ANOVA followed by Bonferroni *post hoc* comparisons for all pairings, * *p* < 0.05, ** *p* < 0.01. Sample sizes are indicated in each bar. Intercrossing GluN1KD and WAVE-Tg mice did not result in improved performance in the puzzle box, open field, or social approach behavior tests.

#### Open field locomotion

To help us understand the effects WAVE-1 rescue might have on other domains of cognitive function, we performed the open field test to assess for differences in locomotor activity and habituation in GluN1KD-WAVE hybrids compared to WT, WAVE-Tg, and GluN1KD littermates. Two-way repeated measures ANOVA reveals a significant effect of genotype on locomotor activity over time in a novel environment (*F_3,83_* = 53.59, *p* < 0.01). As shown in Figure 7b, both GluN1KD and GluN1KD-WAVE mice show increased locomotion and decreased habituation. All pairwise comparisons between a GluN1KD genotype and a non-GluN1KD genotype are significant (Bonferroni adjusted *p* < 0.01), but not between GluN1KD and GluN1KD-WAVE mice. No sex differences were observed in this behavior test. Intercrossing GluN1KD and WAVE-Tg mice does not significantly change GluN1KD’s effects on hyperactivity and habituation as assessed by the open field test.

#### Social approach behavior test

We assessed social approach behavior in WT, WAVE-Tg, GluN1KD, and GluN1KD-WAVE mice to determine the effects of WAVE-1 rescue on an aspect of social cognition. Social approach behavior is quantified as the amount of time a mouse spends in a zone with a novel mouse (30). One-way ANOVA reports a significant effect of genotype on social approach behavior (*F_3,92_* = 5.69, *p* < 0.01). As shown in Figure 7c, GluN1KD mice (202.2 ± 12.4 s) and GluN1KD-WAVE mice (213.1 ± 15.9 s) spend significantly less time in the social zone compared to WAVE-Tg mice (269.9 ± 14.0 s, Bonferroni adjusted *p* < 0.01 vs GluN1KD, *p* = 0.01 vs GluN1KD-WAVE). WT mice similarly spend significantly more time in the social zone (258.7 ± 13.7 s) compared to GluN1KD mice (*p* = 0.04), though it was a trend towards significance when compared with GluN1KD-WAVE mice (*p* = 0.15). Nonsocial zone time, a measure of general object novelty, also showed a significant effect of genotype (*F_3,92_* = 16.84, *p* < 0.01). WT mice spent more time exploring the empty-cage zones compared to GluN1KD mice (141.6 ± 9.3 s vs 63.2 ± 7.7 s, *p* < 0.01) and to GluN1KD-WAVE hybrids (83.7 ± 7.7 s, *p* < 0.01). No sex differences were observed in this behavior test. Social approach behavior is not rescued in GluN1KD-WAVE mice.

### Hippocampal WAVE-1 levels and CA1 pyramidal neuron dendritic spine densities are unchanged across genotypes

Since maze exploration performance is improved in GluN1KD-WAVE mice, we asked whether this effect was due in part to WAVE-1 increases in the hippocampus, key for learning and memory formation (38,39). Hippocampal WAVE-1 expression was measured by western blot, and there were no differences detected between the four genotypes (Figure 8a). (One-way ANOVA *F_3,20_* = 2.33, *p* = 0.11, *β* = 0.50).

**Figure 8.**
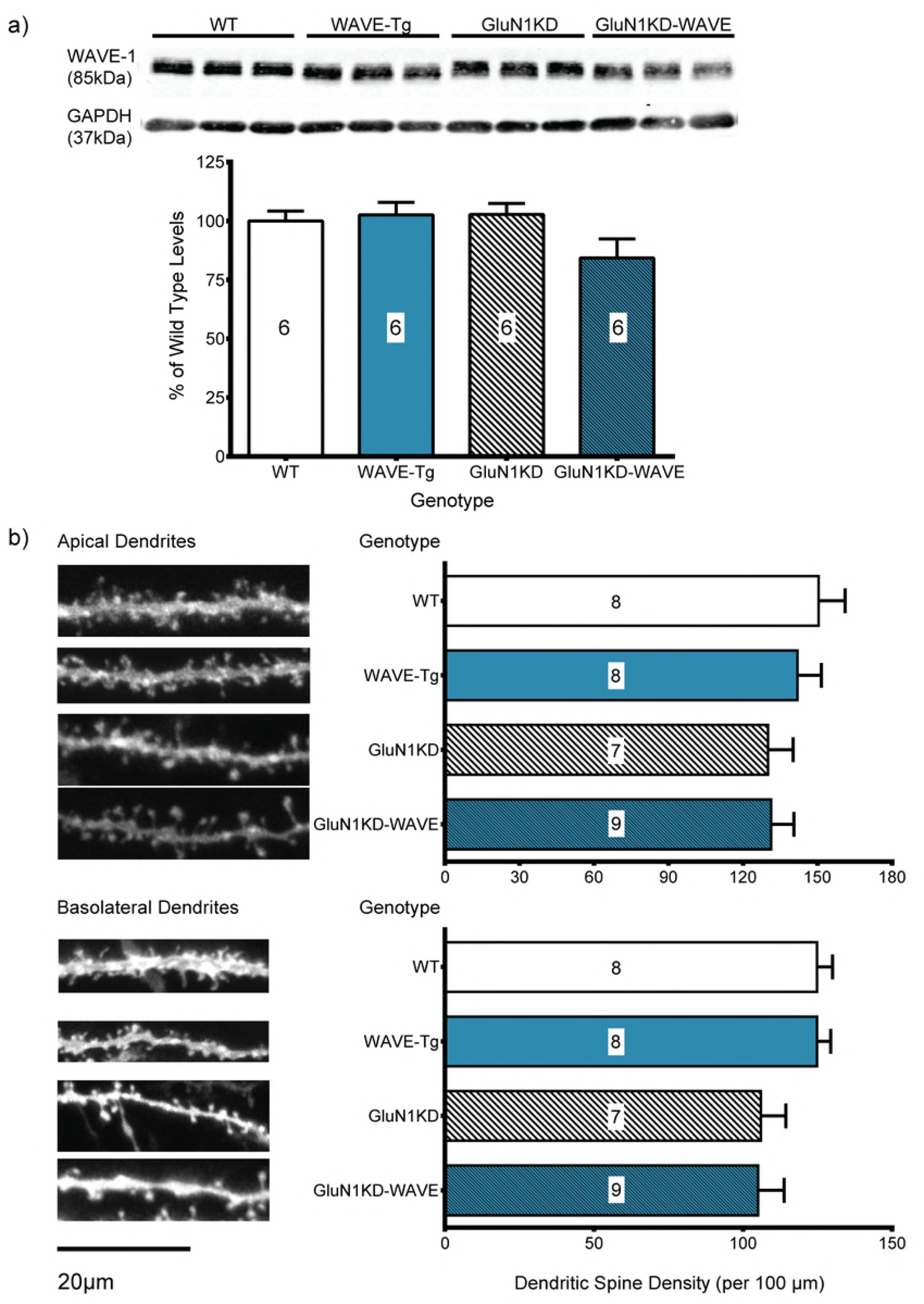
(a) Hippocampal WAVE-1 western blots and (b) CA1 pyramidal neuron spine density analyses in WT, WAVE-Tg, GluN1KD and GluN1KD-WAVE mice. a) Representative western blots are shown on top with their quantifications below. Compared to WT mice, WAVE-1 was not significantly changed in WAVE-Tg, GluN1KD, or GluN1KD-WAVE mice. All blots were first normalized to GAPDH loading control bands before analysis. A sample size of 6 mice was used for every genotype. Data was analyzed by one-way ANOVA followed by Bonferroni *post hoc* comparisons for all pairings. b) Apical (top) and basolateral (bottom) dendritic spine analyses are shown with representative dendrite images on the left and spine density quantifications on the right. GluN1KD mice had less apical and basolateral spines compared to non-GluN1KD mice, but these differences were not significant. Mouse sample sizes are denoted within each bar. 3-6 dendrite sample images were analyzed for each mouse. Scale bars represents 20µm. Data was analyzed by one-way ANOVAs followed by Bonferroni *post hoc* comparisons for all pairings.

Our assessments of CA1 pyramidal neuron dendritic spines also indicated no significant effect of genotype on either apical or basolateral spine density by one-way ANOVAs (*F_3,28_* = 0.93, *p* = 0.44, *β* = 0.77; *F_3,28_* = 2.48, *p* = 0.08, *β* = 0.45; respectively). Representative dendrite images and spine density quantifications are shown in Figure 8b. For apical dendritic spines, GluN1KD mice have a spine density of 130 ± 10 spines/100 μm while GluN1KD-WAVE mice have a density of 131 ± 9 spines/100 μm. These values are lower, but not significantly so, compared to WT levels of 150 ± 11 spines/100 μm and WAVE-Tg levels of 142 ± 10 spines/100 μm. The results are similar for basolateral dendritic spines: GluN1KD mice have 106 ± 8 spines/100 μm while GluN1KD-WAVE mice have 105 ± 9 spines/100 μm, neither significantly different compared to WT levels of 125 ± 5 spines/100 μm or WAVE-Tg levels of 125 ± 5 spines/100 μm. Overall, we did not observe a significant reduction in WAVE-1 or a spine density deficit in the CA1 region of GluN1KD mice. We also did not observe alterations in hippocampal WAVE-1 levels or spine density in WAVE-Tg mice. However, these experiments were determined to be underpowered *post hoc*.

## Discussion

### Synaptic and molecular deficits in GluN1KD mice

Our study identifies a link between NMDAR hypofunction and Rho GTPase signaling that can contribute to age-related spine loss in the striatum. We focused on the integrity of Rho GTPase pathways since they are well-known to regulate dendritic spine architecture (13). After surveying the levels of the principal Rho GTPases and their effectors, we identified two proteins in the same pathway that were altered, Rac1 and WAVE-1.

Changes in these two specific proteins are consistent with a previous report of GluN1KD mice having reduced levels of synaptic DISC1, which is an upstream regulator of Rac1 (7,9,33). Pharmacological blockade of NMDARs is also reported to reduce WAVE-1 in the cortex (40), further suggesting a functional relationship between NMDARs and WAVE-1. WAVE-1 promotes actin polymerization at synapses, affecting synaptic connectivity and cognition (34,35,41). Consistent with our results, knockout of WAVE-1 causes dendritic spine deficits and behavioral abnormalities (16,35). Similar phenotypes are observed in mice with disrupted Arp2/3 complexes that act with WAVE-1 to polymerize actin (42).

Overall, there is strong evidence that a disruption in Rac1 signaling leads to dendritic spine loss and impaired cognition.

Both striatal MSN spine deficits and Rac1 and WAVE-1 deficits are age-dependent. Spine deficits are seen only in older GluN1KD mice while Rac1 and WAVE-1 present a more complex picture of age-dependency. These proteins were also decreased in older mice but vary in younger mice – Rac1 was increased while WAVE-1 was decreased in GluN1KD mice. We previously reported that striatal spine density was unchanged in 2-week old GluN1KD mice (7). In this study, we determined that striatal spine density deficits emerge between 3 and 6 weeks of age. Consistently, the molecular and spine density deficits were seen in ages when most GluN1KD behavioral abnormalities manifest (8). This perhaps also parallels schizophrenia symptom onset in humans (43). Whether the varying molecular and synaptic changes in juvenile GluN1KD mice represent competing compensatory protective mechanisms or initial complex aberrations that ultimately lead to adult behavioral abnormalities is unknown. However, these age-based correlations yet again point to disrupted Rac1 signaling components and dendritic spine abnormalities as being related to GluN1KD behavioral deficits.

### Effect of restoring WAVE-1 on GluN1KD phenotypes

A BAC transgene bearing the human *Wasf1* locus was successfully incorporated into the mice of our study and expresses throughout the brain. Although the transgene increased Wasf1 message levels in both the striatum and hippocampus, the protein levels of WAVE-1 were increased in the striatum, but not in the CA1 region of the hippocampus. This increase in the striatum was sufficient to normalize the WAVE-1 levels in GluN1KD-WAVE mice towards WT levels. Our results suggest that WAVE-1 protein levels may be dictated chiefly by translational regulation and/or post-translational processes, at least at the CA1 region of the hippocampus in the context of NMDAR deficiency in mice.

Rescue of WAVE-1 protein levels in the striatum of GluN1KD-WAVE mice led to an increase in MSN spine density. Consistent with our hypothesis, GluN1KD-WAVE mice had a spine density closer to WT than GluN1KD levels, supporting the biological relevance of targeting WAVE-1, and Rho GTPase signaling in general, to reverse GluN1KD phenotypes. The magnitude of the deficit and subsequent rescue is similar to those observed when spine density changes are associated with significant behavioral changes in Pavlovian conditioning and visual or motor learning (12,44,45). Thus, it is reasonable to expect behavioral outcomes from the increase in striatal WAVE-1 levels and spine density.

Our behavioral assessments were consistent with previous reports of GluN1KD deficits (3,4,8). GluN1KD mice displayed increased locomotor activity, decreased habituation, decreased social approach time, longer puzzle box completion time, decreased Y-maze spontaneous alternations, and worse 8-arm radial maze performance (increased WME and decreased ETR) compared to WT littermates. The selective improvement of maze exploration performance in GluN1KD-WAVE mice is particularly interesting. This improvement is seen in both the Y-maze and 8-arm radial maze. There is evidence for the striatum being a key region for cognition and maze exploration performance (46–49). Improvements of both striatal spine density and maze exploration performance in GluN1KD-WAVE mice are thus consistent with these studies. However, this selective set of improvements point to the possibility of similar synaptic and molecular changes in the hippocampus, which is a key structure for cognitive tasks like maze exploration (50,51).

### Hippocampal and striatal differences in GluN1KD-WAVE mice

Despite improvements in maze exploration, GluN1KD-WAVE mice did not show a change from GluN1KD littermates in hippocampal WAVE-1 protein levels or CA1 dendritic spine density. Improvements of maze exploration performance is often interpreted as an improvement of spatial learning and memory (42,48,52), which requires intact hippocampal function (38,39,53). The specific improvement of maze exploration performance in GluN1KD-WAVE mice, independent of changes in hippocampal WAVE-1 or CA1 pyramidal neuron spine density, is therefore unexpected. It should be noted that spine density measures of hippocampal neurons were found to be underpowered *post hoc* (*β* = 0.45-77). New sample-size calculations using effect-size numbers based on our data yielded a required n of 10 – 33 per group for 1- *β* = 0.8, greatly exceeding typical sample sizes for dendritic spine studies (23,35,41). However, if our results were indeed false negatives, we would predict CA1 spine density deficits in the hippocampus of GluN1KD mice but not rescue in GluN1KD-WAVE mice. The improvement of maze exploration performance in GluN1KD-WAVE mice therefore still seems incongruent with our data showing no hippocampal WAVE-1 or CA1 spine density rescue.

Multiple avenues can be taken to further investigate this discrepancy between our hippocampal and behavioral data. For instance, there is the need to consider WAVE-1 activation in GluN1KD and GluN1KD-WAVE mice. Changes in NMDAR levels may have more effects on the activity of WAVE-1 than on the levels of total WAVE-1 at the hippocampus, unlike what we have seen in the striatum. WAVE-1 phosphorylation may be one useful indicator, and is generally considered to be inhibitory (41,54,55). It is also possible that dendritic spines aside from those of CA1 pyramidal neurons in the hippocampus could be key for the rescue seen in GluN1KD-WAVE mice, such as those in the CA3 or dentate gyrus regions – also key for memory and maze exploration tests (56,57). Finally, a lack of physical changes in mouse CA1 pyramidal neuron spines has been reported before in neurons with eliminated AMPA and NMDA receptor signaling *in vivo* (58). Instead, the authors found drastically altered electrical and functional characteristics in these neurons. This could be applicable to our GluN1KD mice, independent of the *Wasf1* transgene. There is still much to study regarding the molecular and synaptic underpinnings of maze performance improvements in GluN1KD-WAVE mice.

Despite the lack of observed hippocampal rescue in GluN1KD-WAVE mice, our striatal data provides important clues about the state of cognition in these hybrids. In previous studies, the dorsal striatum was linked to reinforcement of stimulus-response associations (46), the nucleus accumbens was linked to goal-directed exploration (47), and the marginal division of the striatum was linked to both early and late stage long-term memory consolidation (49). We did not use overt stimuli or cues in either of our maze tests, while goal directed exploration and long-term memory were important factors in our repeated 8-arm maze trials of food-restricted mice. The molecular and synaptic rescue we observed at the striatum of GluN1KD-WAVE mice may thus have influenced their learning of food-foraging strategy. Similarly, the improved Y-maze performance of GluN1KD-WAVE mice may be due to it having fewer arms and being analogous to non-delayed random foraging tasks, which also depended on the nucleus accumbens (47). However, neither the nucleus accumbens nor the marginal division of the striatum were linked to cognition independently of the hippocampus (47,49). Thus, the hippocampus of GluN1KD-WAVE mice may be rescued in a manner we did not assess or the effects of the *Wasf1* transgene at the striatum and elsewere were sufficient to have a noticeable effect on maze exploration performance.

With our current data of GluN1KD-WAVE mice, we see striatal WAVE-1 and MSN spine density rescue along with a specific behavioral rescue in Y-maze and 8-arm radial maze performance. Such a rescue may be related to long-term memory and goal-directed exploration. To note, maze tests have been interpreted as evaluations of various aspects of cognition, such as habituation, curiosity, spatial working memory, and more (59). Studies making such interpretations sometimes do so independently of concurrent assessments of molecular or cellular changes, limiting the interpretability of behavioral data. There is a need of more research and clearer definitions of the relationships between specific brain changes, behavioral changes, and their interpretations as cognitive aspects.

## Acknowledgements

The authors would like to acknowledge and thank Wendy Horsfall for help with animal husbandry and taking care of the animals used in this study and Gloria Hu for preliminary work on striatal dendritic spine analysis.

